# Direct causality measures unravel complex networks of cardiovascular oscillations and their modifications with postural stress

**DOI:** 10.1101/2025.03.26.645432

**Authors:** Chiara Barà, Laura Sparacino, Luca Faes, Michal Javorka

## Abstract

This study provides a comprehensive investigation of the spontaneous short-term regulatory mechanisms affecting cardiovascular and cardiorespiratory interactions during supine rest and in response to postural stress. The direct causality measure of conditional transfer entropy was applied to beat-to-beat heart period, arterial pressure, respiration, and arterial compliance variability series assessed in thirty-nine healthy subjects during the supine resting state and the orthostatic challenge. The inferred physiological networks behind these two conditions reveal well-known regulatory mechanisms, such as the tilt-induced decreased respiratory sinus arrhythmia (RSA) and increased baroreflex, as well as less explored interactions such as those involving compliance, which suggest striking physiological responses. Specifically, we found tight relationships between compliance and heart period, arterial pressure and respiration, which advocate the non negligible involvement of this cardiovascular parameter into the intricate hank of the most studied physiological interconnections. Furthermore, the joint use of parametric and model-free estimation approaches allowed us to infer the prevalence of linear and nonlinear dynamics, as well as the effects on the inferred directed links of low- and high-frequency oscillations reflecting autonomic modulation. In conclusion, our study proves that direct causality measures are crucial to assess the characteristic links of complex cardiovascular networks and infer the many underlying short-term regulatory mechanisms.

**NEW & NOTEWORHTY:** While short-term regulatory mechanisms involving heart period, respiration and arterial pressure have been widely investigated, the way they produce and buffer cardiovascular oscillations in different physiological states is not fully understood. This study proposes a thorough investigation of a four-node physiological network, including the less explored arterial compliance variability, and provides insights into the linear vs. nonlinear characterization and spectral content of the causal dynamics representing each link within the network.

## INTRODUCTION

The human organism can be seen as an integrated network constituted by multiple organ systems, each characterized by its own structural organization and functional complexity, which continuously interact to coordinate their functions, generate distinct physiological states in response to internal, external and pathological perturbations, and maintain homeostasis [1, 2]. In the field of Network Physiology, which focuses on the coordination and network interactions among diverse biological systems and subsystems, data-driven methods for network inference can be exploited to build network models from sets of observed multivariate time series describing the activity of the network nodes [2, 3]. Such models are usually represented as graphs, where nodes represent different physiological sub-systems or districts, and links map functional dependencies between these systems. Straightforward examples involve the well-known cardiovascular interactions between the heart rate and arterial pressure variabilities [4], as well as the tangled coordination between the cardiac and the respiratory subsystems [5]. Cardiovascular and cardiorespiratory networks, which reflect the modulation of heart rate, arterial pressure and respiratory variabilities [6], have been largely studied, with the aim of disentangling both autonomic regulation mechanisms and mechanical effects occurring in diverse physiological states and conditions [7, 8].

As physiological networks depend on a big variety of mechanisms, an appropriate characterization of their dynamics would require the involvement of many variables: cardiovascular and cardiorespiratory loops represent only a small portion of a more complex and wider system. While heart rate, arterial pressure and respiration still remain the most studied [6], advancements in the beat-to-beat analysis of physiological variables have recently emerged. For instance, little is known about the short-term-variability nature of arterial compliance, a cardiovascular variable characterizing mechanical and structural properties of the arteries [9, 10]. Arterial compliance variability is expected to be directly affected by the sympathetic nervous system, and mostly indirectly by vagal activity in different patho-physiological conditions, as well as by heart rate, blood pressure and respiratory variabilities [10–12]. Investigating how this parameter behaves within complex physiological networks is essential to characterize its dynamics for physiological research and clinical purposes. Furthermore, physiological mechanisms are challenged by a number of stressors, e.g., postural stress induces a reorganization of cardiovascular oscillations and of their coupling related to the shift in the sympatho-vagal balance towards sympathetic activation and parasympathetic withdrawal [13, 14]. Hence, probing the investigated network after its modification due to a given stressor is of remarkable importance to characterize the type and modalities of network adaptation.

From a methodological viewpoint, noninvasive analysis of spontaneous oscillations of physiological variables has typically been performed by multivariate time series analysis. Classical linear and nonlinear approaches defined in the time, frequency and information-theoretic domains, such as those based on cross-correlation analysis [15], nonlinear prediction [16, 17], coherence analysis [15, 18], and entropy measures [6, 19], have been used to describe the system properties captured by network models. However, these methods are overall limited by their intrinsic pairwise formulation (i.e., only two network nodes are taken into account), thus neglecting potential confounding effects due to non-involved variables [20, 21]. Moreover, they are generally non causal (i.e., measures are symmetric and non directed), meaning that they do not allow identification of the direction of the information flow between the considered processes. Nevertheless, understanding driver–response relationships between physiological sub-systems from the analysis of spontaneous cardiovascular oscillations is of utmost importance, since it allows a noninvasive, detailed comprehension of physiological regulatory mechanisms [22]. In this context, past studies have been focused only on specific network links, such as the cardiovascular or cardiorespiratory closed-loops with feed-forward and feedback mechanisms describing the pair-wise interplay between the heart and vascular or respiratory systems, respectively [6, 19, 22]. Indeed, various time series analysis techniques have been developed for the quantification of these specific physiological coupling mechanisms, e.g., the baroreflex (BR) sensitivity as a measure of reflex autonomic control of the cardiovascular system [18] or the amplitude of respiratory sinus arrhythmia (RSA) as an index of cardiac vagal tone [22]. Whilst recent efforts have been oriented to capture the complex dynamics involving three or more processes [19, 21, 23–25], studies that investigate and interpret the role of specific causal links among multiple physiological nodes in an exhaustive way are limited in the analysis of cardiovascular oscillations. Among these approaches, methodologies exploiting conditional causality approaches in multivariate contexts would allow to estimate the strength of the directed coupling between two given nodes canceling out the influence of the rest of the network.

In this context, the aim of the present work is to elicit noninvasively the physiological mechanisms underlying complex cardiovascular regulation from the joint analysis of the spontaneous beat-to-beat variability of the main cardiovascular and respiratory parameters. To this end, we carry out a systematic analysis of direct causal interactions in a physiological network comprising four variables, i.e., heart period, arterial pressure and respiration, including also the rarely explored arterial compliance beat-to-beat time series. We exploit diverse methodologies, namely parametric [26] and non-parametric [27, 28] conditional causality approaches designed to disentangle putative linear and nonlinear dynamics, respectively. Furthermore, parametric spectral analysis provides valuable insights into the frequency domain of physiological time series displaying a rich oscillatory content, as in the case of cardiovascular and respiratory variables [29, 30]. We reconstruct the structure of the investigated network in terms of presence/absence of direct links between pairs of variables, in the supine resting state and in response to the homeostatic alterations caused by the postural stress. The physiological relevance and meaning of each of the observed interactions is first statistically validated and then accurately discussed to either confirm or confute previous results, but also to provide novel insights on the complex mechanisms governing the cardiovascular, cardiorespiratory and vascular-respiratory interplays.

## I. MATERIALS AND METHODS

### A. Study protocol

This study included 39 young and healthy Caucasians (22 women, 17 men; median age: 18.7 yr), and was approved by the ethical committee of the Jessenius Faculty of Medicine, Comenius University of Bratislava [10, 31– 33]. All subjects or their legal representatives, in participants under 18 years of age, gave written informed consent before the examination. Subjects were positioned on a motorized head-up tilt table with a foot support, and were asked to avoid unnecessary speaking and moving during the measurement. The study protocol consisted of two consecutive phases: supine rest (REST, 15 min), and head-up tilt (HUT, the subject was tilted to 45 degrees for 8 min to evoke mild orthostatic stress). There were no signs of presyncope in any subject during the orthostatic challenge. All subjects breathed spontaneously without any effort to control breathing rate or tidal volume.

### B. Data acquisition

Electrocardiogram (ECG, CardioFax ECG-9620, NihonKohden Japan), and arterial blood pressure (BP) curve from finger, with the subsequent brachial arterial BP reconstruction by the photoplethysmographic volume-clamp method (Finometer Pro, FMS Netherlands), were simultaneously and noninvasively recorded (Fig 1a, ECG and BP). A hydrostatic height correction unit was attached on the measured arm at the level of heart to correct for effects of hydrostatic pressure associated with the vertical distance between finger and heart level. Respiration signal (Breath) was measured through the respiratory inductance plethysmography method (RespiTrace, NIMS, Miami Beach, FL, USA) using both thoracic and abdominal impedance belts following qualitative diagnostic and fixed volume calibrations (Fig 1a, Breath). Impedance cardiography (ICG, CardioScreen 2000, Medis, Germany) was also performed, thus enabling continuous beat-to-beat noninvasive monitoring of several indices characterizing myocardial performance and hemodynamics. This method calculates first the changes in the transthoracic impedance (IMP or Δ*Z* waveform) as a result of the volume and blood flow velocity variations in the aorta, and then the changes in blood volume in the transthoracic region over time, getting the derived IMP waveform as the first mathematical derivative of Δ*Z* waveform, indicated as *dZ/dt* (Fig 1a, *dZ/dt*). All the acquired signals were digitized at a sampling rate of 1 kHz.

**Figure 1.**
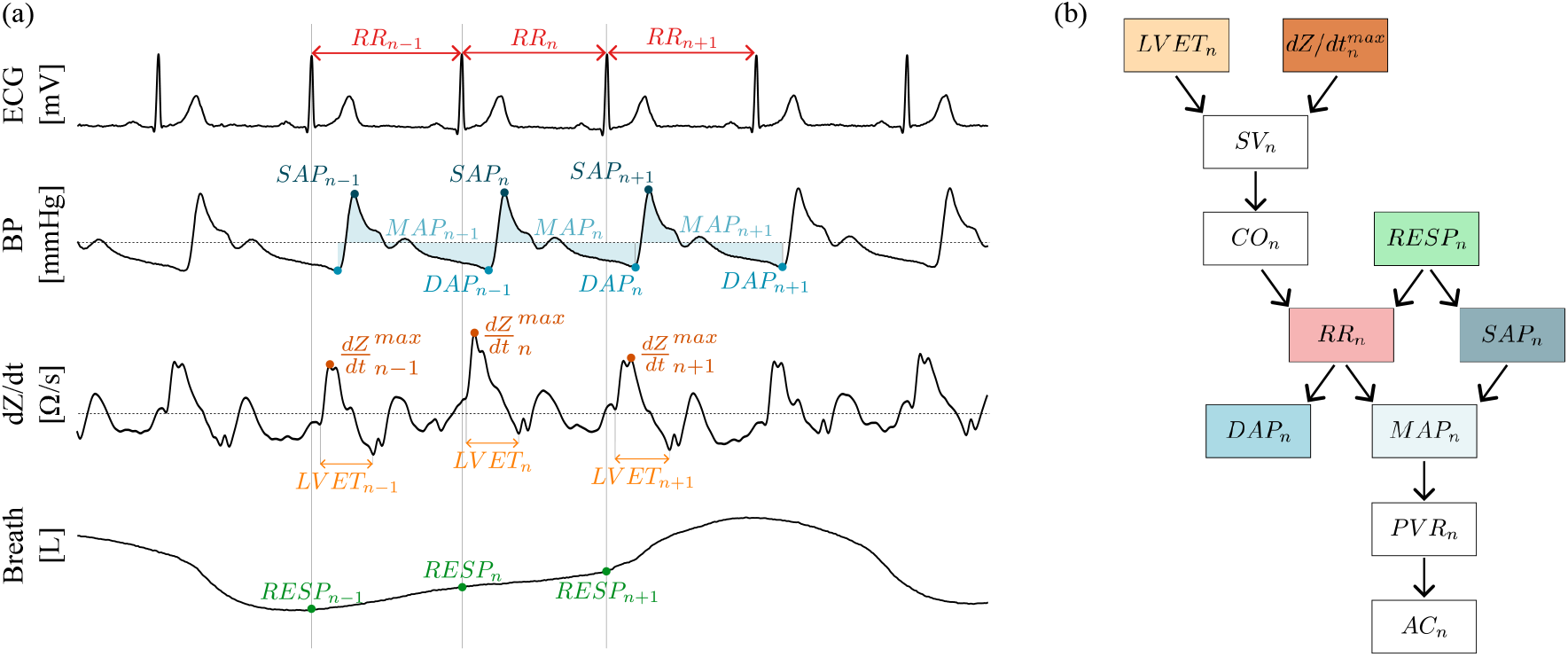
**(a)** Acquired signals (electrocardiogram, ECG; blood pressure, BP; impedance cardiography, *dZ/dt*; respiration signal, Breath) and corresponding extracted time series (heart period, RR; mean arterial pressure, MAP; respiration amplitude, RESP). Arterial compliance (AC) was computed exploiting parameters extracted from the *dZ/dt* and the BP waveforms. **(b)** Convention for the choice of zero-lag effects; for visual purposes, colors of the boxes correspond to the exemplary time series points depicted in panel a.

### C. Time series extraction and data pre-processing

Starting from the acquired signals, physiological time series displaying the dynamic activity of cardiovascular and respiratory variabilities were extracted, as detailed in Fig. 1a. Heart period (RR) intervals were approximated as the time distance between the *n*^*th*^ and the (*n*+1)^*th*^ R peaks of the ECG (i.e., RR_*n*_); the *n*^*th*^ systolic arterial pressure (SAP) value (i.e., SAP_*n*_) was measured as the maximum of the BP signal inside RR_*n*_. The *n*^*th*^ diastolic arterial pressure (DAP) value (i.e., DAP_*n*_) was taken as the minimum of BP between the occurrences of SAP_*n*_ and SAP_*n*+1_. Mean arterial pressure (MAP) was calculated as the true integrated mean pressure between the occurrences of DAP_*n*−1_ and DAP_*n*_. The *n*^*th*^ respiration amplitude (RESP) value (i.e., RESP_*n*_) was computed sampling the Breath signal on the *n*^*th*^ R peak of the ECG.

Cardiovascular measures including stroke volume (SV) and cardiac output (CO) were estimated starting from the *dZ/dt* waveform. Specifically, the Bernstein and Sramek formula [34] was exploited to compute SV, which is proportional to the maximal systolic ejection speed (*dZ/dt*^*max*^) and to the duration of the ejection phase (Left Ventricular Ejection Time, LVET) evaluated in the same beat, while the *n*^*th*^ CO value (i.e., CO_*n*_) was computed as the ratio between SV_*n*_ and RR_*n*−1_ [34, 35]. Peripheral vascular resistance (PVR) was calculated for each heartbeat (i.e., PVR_*n*_) as the ratio of MAP_*n*_ and CO_*n*_, assuming zero venous pressure at the right atrium. Finally, the value of arterial compliance (AC) was quantified on a beat-to-beat basis through a recently developed method, based on a reliable estimation of the time constant *τ*, i.e., the rate of the peripheral BP decay during the diastolic phase, as well as on the exploitation of the common relationship between *τ*, AC and PVR based on the two-element Windkessel model, i.e., *AC*_*n*_ = *τ*_*n*_*/PV R*_*n*_ [9].

As shown in Fig. 2a for a representative subject, stationary segments of 300 consecutive beats were extracted from the original RR, MAP, AC, and RESP time series (henceforth referred to as *H, M, C*, and *R*, respectively). These windows were selected in REST starting 8 min after the beginning of the measurement, and in HUT 3 min after the position change from supine to tilt, in order to avoid transient changes in cardiovascular parameters. We refer the reader to [10, 31–33] for further details about the study protocol, data acquisition and time series extraction.

**Figure 2.**
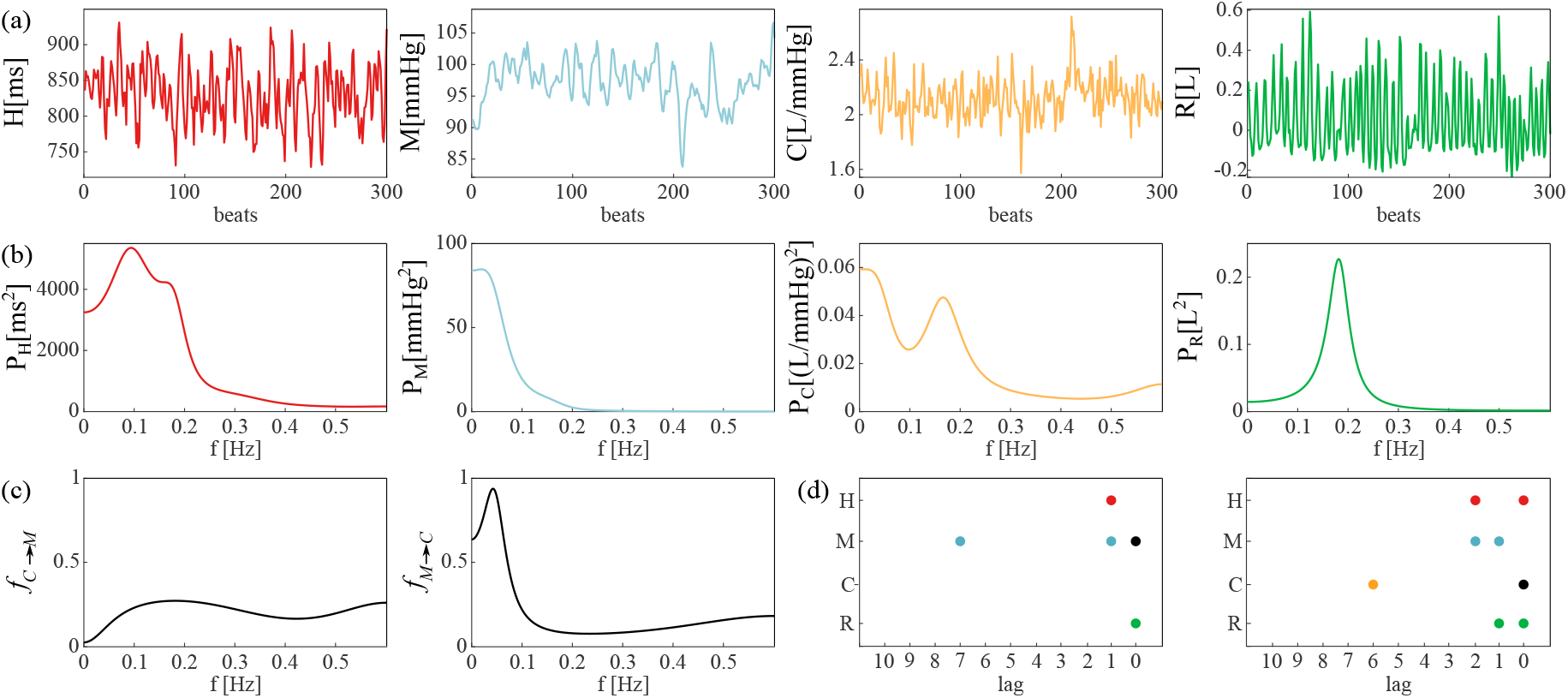
**(a)** Time series *H* (RR), *M* (MAP), *C* (AC), *R* (RESP), and **(b)** their parametric spectra computed for a representative subject in the supine resting state. Exemplary pair of variables (*C, M*): **(c)** spectral profiles of the conditional causality measures obtained through the frequency-domain model-based (MB) approach (*f*_*C*→*M*_, *f*_*M* →*C*_) [29], and **(d)** lag selection for the computation of non-uniform embedded conditional transfer entropies through the model-free (MF) approach (*T*_*C*→*M*_, *T*_*M* →*C*_).

Classical time domain markers, i.e., the mean and standard deviation of RR (*µ*_*H*_ [*ms*] and *σ*_*H*_ [*ms*]), MAP (*µ*_M_ [*mmHg*] and *σ*_M_ [*mmHg*]), and AC(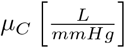 and 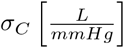), as well as the respiratory rate and tidal volume (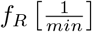 and 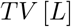), were first computed. Then, time series were pre-processed to remove slow trends through high-pass filtering (zero phase, cut-off frequency 0.0156 Hz) and normalized to zero mean.

### D. Conditional causality measures

In this work, physiological interactions at rest and in response to orthostatic stress were investigated through the information-theoretic measure of conditional/partial transfer entropy (TE) proposed and discussed in [36], with the purpose to identify and quantify the direct causal links among the *H, M, C*, and *R* time series constituting the four nodes of the investigated network. This measure expands the concept of bivariate TE [37], canceling out the influence of the rest of the network on estimating the strength of directed (i.e., causal, time-asymmetric) coupling between two given nodes. Thus, it resolves whether the interaction between the two considered dynamic random processes is direct or mediated by the other processes.

Specifically, two different formulations of the conditional TE were here applied to physiological data, i.e., the autoregressive (AR) model-based (MB) [26, 29, 30, 38, 39] and model-free (MF) [27, 28, 40, 41] formulations.

#### 1. Model-based formulation of conditional causality

Assuming that the oscillatory activity of the network nodes can be described by means of linear Gaussian stationary processes, the measure of causality defined via TE is equivalent to the concept of Wiener-Granger causality (GC) [42]. In this frame, time [26] and frequency [29] domain formulations of conditional GC based on multivariate linear AR models can be used to characterize the causal relationships among the multiple nodes of the investigated physiological network. Model orders (i.e., maximum lags used to describe lagged interactions within and between processes) were selected for each subject according to the multivariate version of the Akaike Information Criterion (AIC), while model identification was performed using the vector least-squares method [43]. First, time domain versions of conditional GC were computed from the estimated AR model parameters following the procedure described in [26] and detailed in the Appendix (Sect. A.1.1). Remarkably, instantaneous effects involving zero-lag interactions were included in these models. Then, parametric spectra were computed from the estimated AR model parameters by setting the sampling frequency *f*_*s*_ equal to the inverse of the mean RR for each subject. Spectral profiles of the conditional GC measures were thus obtained following the procedure described in [29, 30] and detailed in the Appendix (Sect. A.1.2). Importantly, instantaneous effects were not included in these models.

In Fig. 2b, we show the parametric power spectral densities of the *H, M, C*, and *R* processes for a representative subject in the supine resting state. The spectral profiles of the two conditional GC measures computed for the pair {*C, M*} are depicted in Fig. 2c, showing how the causal information flow from mean BP to arterial compliance (*f*_*M*→*C*_) is located at the very low frequencies (i.e., below 0.1 Hz) for this representative subject. Remarkably, integration of these profiles within the low frequency (LF, *f* ∈ [0.04 − 0.15] Hz) and the high frequency (HF, *f* ∈ [0.15 − 0.4] Hz) bands of the spectrum was performed to get the corresponding LF and HF measures of conditional GC, respectively.

#### 2. Model-free formulation of conditional causality

The most widely used continuous MF estimator, i.e., the *k*-nearest neighbour (KNN) method, was applied to capture putative nonlinear behaviours often attributed to physiological mechanisms. Based on searching neighbor samples to approximate the probability distribution of the observed variables [40], the Kraskov-Stögbauer-Grassberger (KSG) formulation presented in [41] was used here to quantify the causal information transferred among the nodes of the network, as previously done in [28, 44] by using the bivariate TE and in [27, 28] by using the conditional TE. In contrast to the linear method, one of the limitations of the MF estimator lies in its sensitivity to the length of the time series and of signal patterns used for the analysis [45]. To overcome this issue, the non-uniform embedding technique described in [28] was exploited to select the most informative time-lagged samples in evaluating the information exchanged across the interacting processes.

In Fig. 2d, the present state of the target process (black dot) and the selected lags of all the time series are reported for the exemplary case of the conditional TE computed along the directions *C* → *M* (left panel) and *M* → *C* (right panel), for a representative subject in the supine resting state. Further details on the MF analysis are reported in the Appendix of this manuscript (Sect. A.2).

As regards data analysis, time series were pre-processed as described in Sect. I C and normalized to have unit variance. Then, non-uniform embedding was applied by fixing a maximum lag of 10 samples, while a number of neighbors *k* = 10 was set to estimate the time domain causality measures.

#### 3. Instantaneous effects among physiological signals

The assessment of causality, intended as the influence that a driver process exerts on a given target, requires a proper handling of instantaneous (zero-lag) effects. In practical physiological time series modelling, instantaneous causality shows up whenever the time resolution of the measurements is lower than the time scale of the lagged causal influences occurring among the analysed processes. This non-delayed effect can arise due to non-physiological factors (e.g., unobserved confounders) or fast (i.e., within-beat) physiologically meaningful interactions [46, 47]. The importance of considering instantaneous effects in the analysis of cardiovascular interactions, where zero-lag interdependencies are expected to contribute significantly to the BR mechanism [46], and of cardiorespiratory interactions, where the information transfer from respiration to heart rate variability is expected to be negligible in the LF band [48], was previously documented [26, 46].

In this study, zero-lag effects were suitably determined according to the measurement convention displayed in Fig. 1a. This was done by visually inspecting the temporal sequence of the investigated physiological indices or the parameters directly involved in their estimation, as well as by considering that the cause always precedes its effect. Specifically, as depicted more in detail in Fig. 1b, zero-lag effects were set along the directions *RESP*_*n*_ → *RR*_*n*_, *RESP*_*n*_ → *MAP*_*n*_, *RESP*_*n*_ → *AC*_*n*_, *RR*_*n*_ → *MAP*_*n*_, *RR*_*n*_ → *AC*_*n*_, and *MAP*_*n*_ → *AC*_*n*_. Remarkably, these zero-lag interactions were selected whenever required by nonparametric (i.e., MF) estimators, and set a priori in the MB extended AR models to compute time domain conditional GC measures. However, the spectral formulation of the MB method could not take instantaneous interactions among signals into account, due to the impossibility to model such interactions in the frequency domain [29]. Neglecting instantaneous influences has a great impact on the proper assessment of frequency-specific linear GC measures [46].

### E. Surrogate and statistical data analysis

The statistical significance of the causality measures estimated by using the MB and MF approaches was assessed for each subject, in both REST and HUT conditions. One-hundred surrogate time series were generated (i) through the *iterative Amplitude Adjusted Fourier Transform* (iAAFT) procedure [49], as regards the MB indices, and (ii) by randomly shifting the target over time (minimum shift of 20 lags) and leaving all the other series unchanged, as regards the MF indices. The MB or MF causality measure computed on the original time series was deemed as statistically significant if its value was higher than the 95^*th*^ percentile of the distribution derived by computing the same measure on the set of surrogates. In this work, the degree of significance *s* for a given MB or MF conditional causality measure, intended as the ratio between the number of subjects for which the measure was deemed as statistically significant over the total number of subjects, was subdivided into three classes, namely *s* = [0 − 50] % (low significance), *s* = [50 − 75] % (medium significance) and *s* = [75 − 100] % (high significance). The first of these classes was arbitrarily intended as not statistically relevant.

Given the small size of the surveyed population, non-parametric statistical tests were applied to assess statistically significant differences between physiological indices evaluated in the two phases of the experimental protocol, i.e., REST and HUT conditions. Specifically, the Wilcoxon signed rank test for paired data was applied on classical time domain markers, i.e., mean and standard deviation for *H, M*, and *C* time series, respiratory rate and tidal volume for *R* time series, and on conditional causality measures evaluated for each pair of nodes using both MB and MF approaches, as well as considering measures integrated in the LF and HF bands of the spectrum. For all the statistical tests, the significance level was set to 0.05.

## II. RESULTS

### A. Classical time domain markers of physiological time series

Fig. 3 shows the results of the analysis performed on classical markers of mean and standard deviation of cardiovascular variability time series, and on respiratory rate and tidal volume as regards the breathing signal. We observe a significant decrease of the mean *H* and *C*, as well as of their variabilities (panels a and c). On the other hand, the significant decrease of *M* is accompanied by a significant increase of its variance (panel b). As regards respiratory indices, a significant decrease of the median values of the respiratory rate and an increase of the tidal volume are found with HUT (panel d).

**Figure 3.**
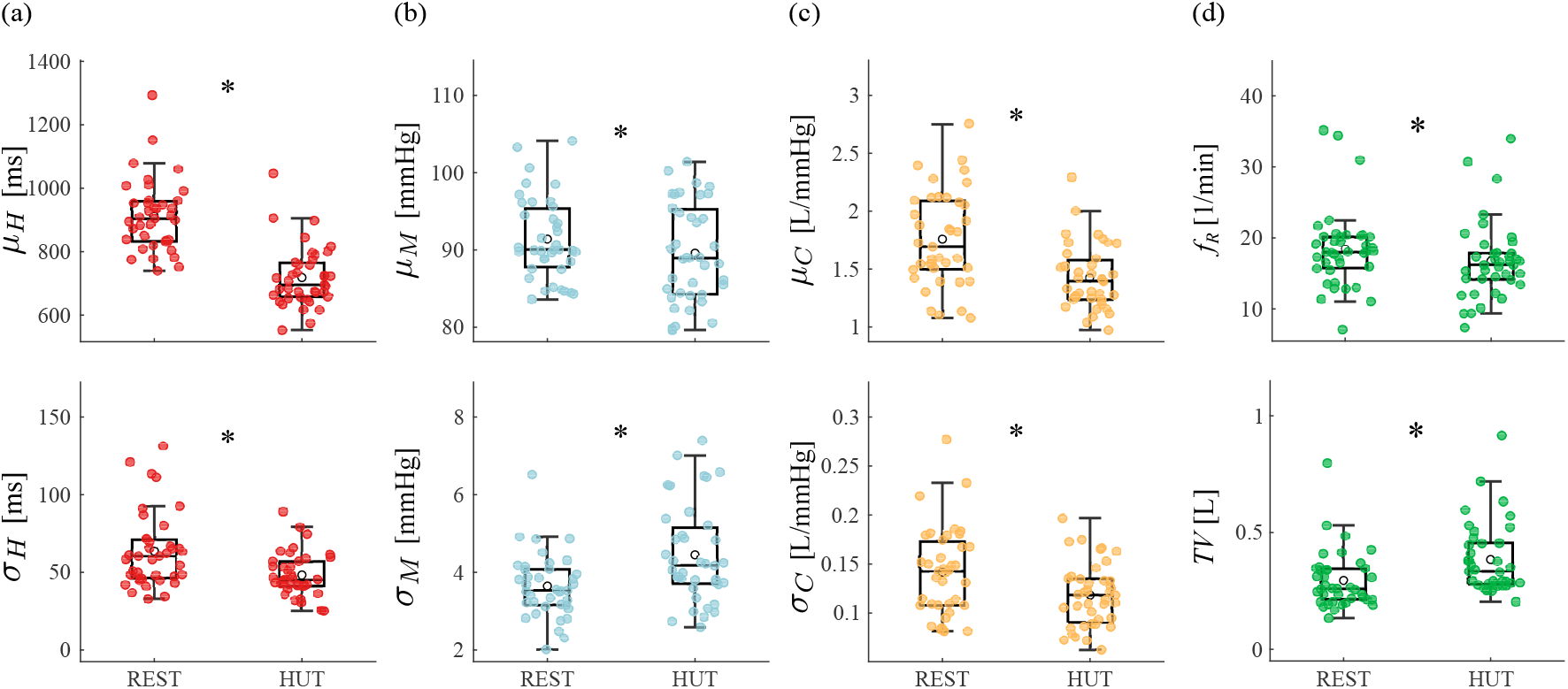
Mean (top row) and standard deviation (bottom row) of the **(a)** *H*, **(b)** *M* and **(c)** *C* time series depicted as boxplot distributions and individual values in the supine (REST) and orthostatic (HUT) positions; **(d)** respiratory rate (top row) and tidal volume (bottom row) computed on the Breath signal. Wilcoxon signed rank test for paired data: REST vs. HUT, (*) *p <* 0.05.

### B. Time domain analysis: model-based vs. model-free approaches

In this section, we display results relevant to Fig. 4, with the aim to compare MB and MF approaches considering the statistical significance of the interactions in the investigated physiological network.

**Figure 4.**
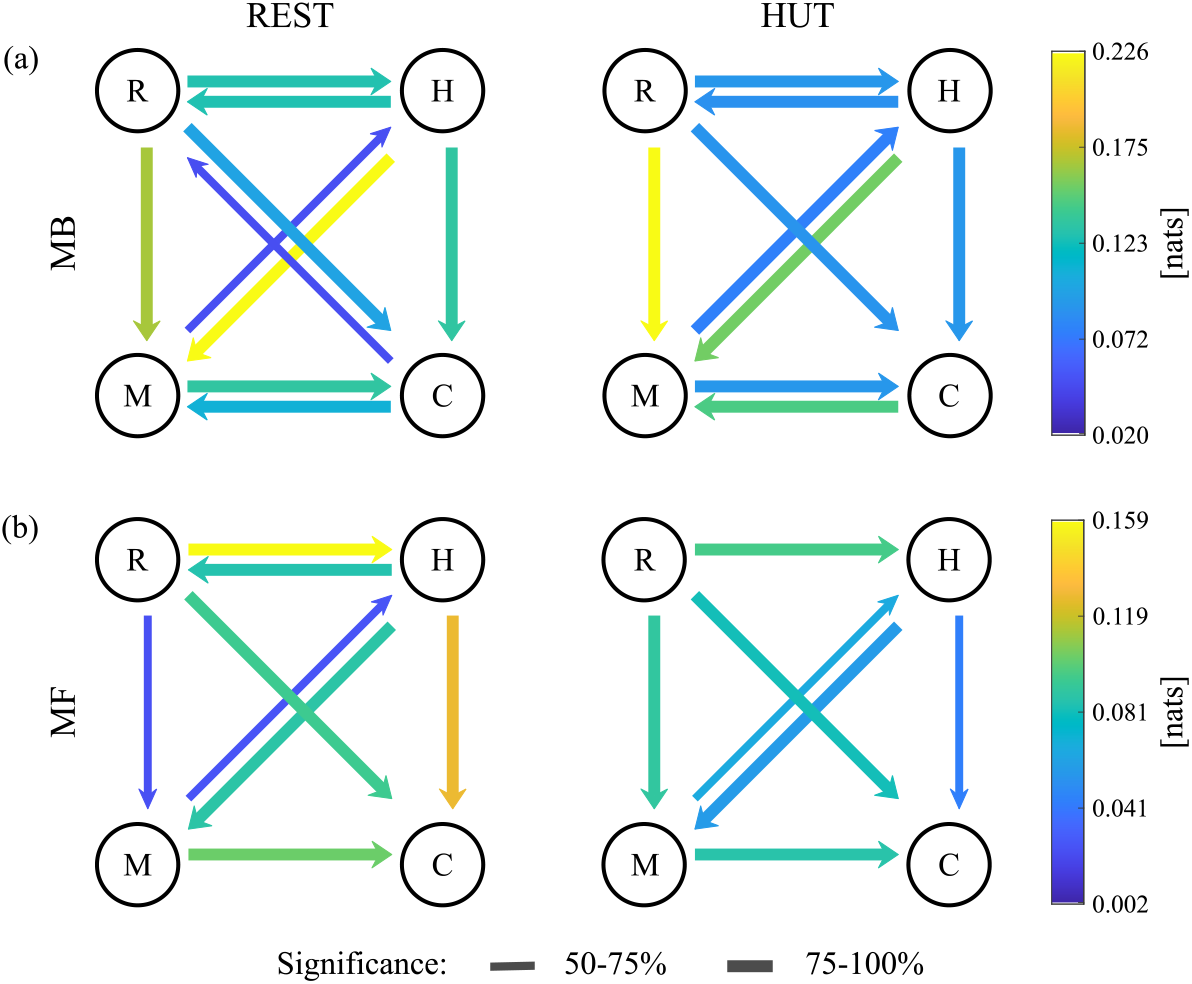
Directional networks computed from **(a)** model-based (MB) and **(b)** model-free (MF) conditional causality measures in the REST (left) and HUT (right) conditions. The strength of the causal couplings within the network is quantified in natural units ([nats]) of information. The different width of the arrows displays results from surrogate data analysis: absence of an arrow indicates low significance (*s <* 50%), a thin arrow indicates medium significance (50% ≤ *s <* 75%), while a thick arrow indicates high significance (*s* ≥ 75%).

Cardiorespiratory dynamics are strongly detected by both MB and MF approaches during REST and HUT states, as shown in Fig. 4a,b. Remarkably, the causal interaction *R* → *H* strongly prevails over the opposite direction if these dynamics are investigated using MF causality measures (Fig. 4b); the latter capture a negligible causal interaction along the direction *H* → *R* during HUT (*s <* 50%). As regards cardiovascular interactions, i.e., the well-known relationship between *M* and *H*, the BR direction (*M* → *H*) is significantly captured in both conditions and by both approaches, with *s >* 75% during HUT using the MB method (Fig. 4a, right) and 50% *< s <* 75% in all the other cases. The opposite link (the so-called feedforward direction, *H* → *M*) seems to represent a stronger mechanism captured by both MB and MF approaches with *s >* 75% (Fig. 4a,b). The directed effect of MAP on AC (*M* → *C*) is always captured by both MB and MF approaches (*s >* 75%), as shown in Fig. 4a,b. The opposite interaction *C* → *M* is significantly detected only by the MB approach (*s >* 75%, Fig. 4a). The causal interaction directed from the heart to large vessels (*H* → *C*) is disclosed by both approaches with high significance, except for the MF measure in the HUT state (50% *< s <* 75%, Fig. 4b, right). The directed effect of respiratory variability on compliance of arteries (*R* → *C*) is always detected with the highest significance by both approaches (*s >* 75%, Fig. 4a,b), while the opposite influence of compliance on respiration seems to have an important role only during the supine resting state if investigated with the MB approach (50% *< s <* 75%, Fig. 4a, left), even though absolute values are very low. The causal interaction *R* → *M* is always observed with the highest significance (*s >* 75%) in HUT using both MB and MF causality measures (Fig. 4a,b, right), and at REST using the MB approach only (Fig. 4a, left). The opposite interaction is not significantly relevant.

### C. Frequency domain analysis: low-frequency vs. high-frequency oscillations

In this section, we display results relevant to Fig. 5, with the aim to compare the low and high frequency characters of the multiple interactions in the physiological network.

**Figure 5.**
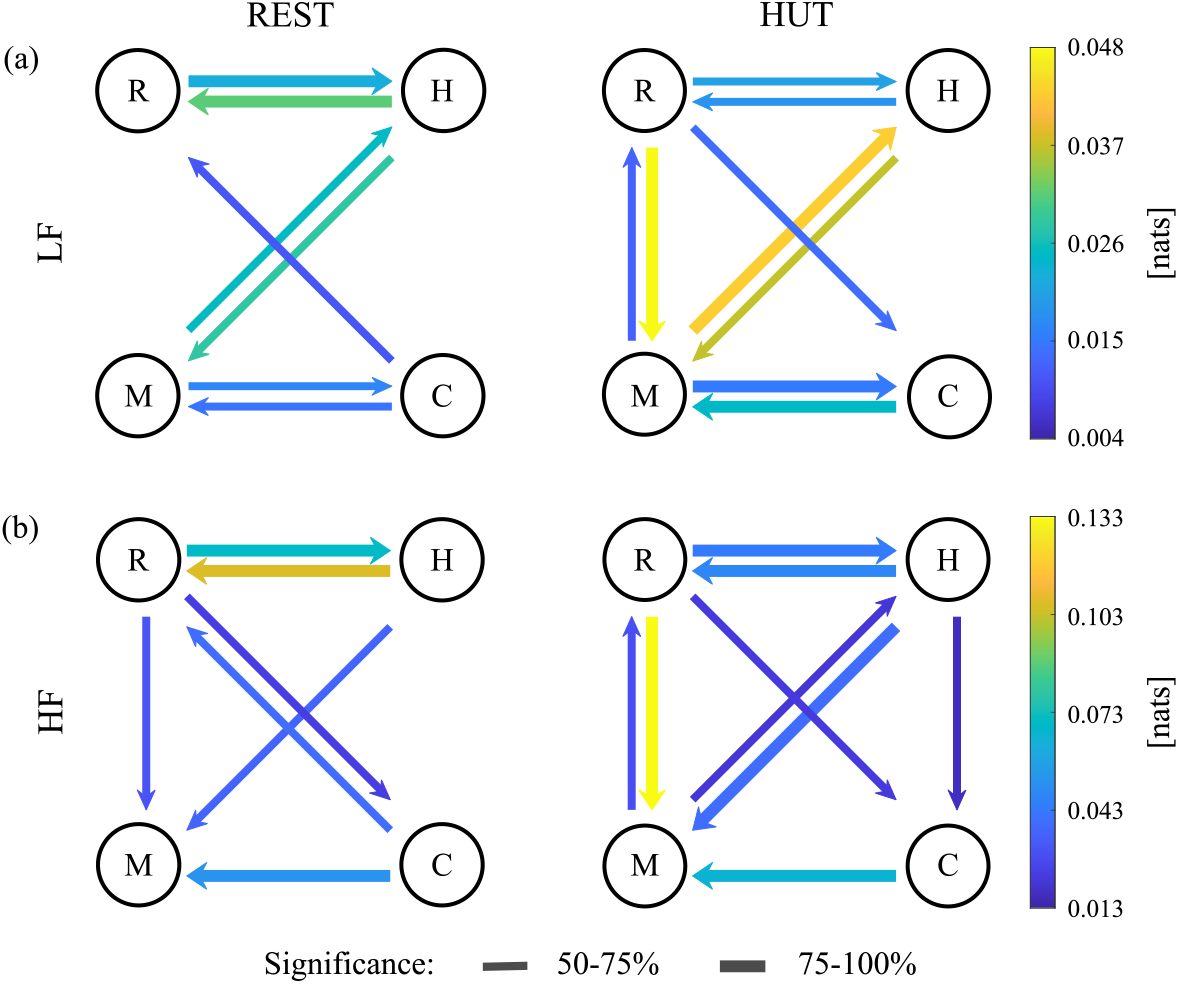
Directional networks computed for conditional causality measures integrated in the **(a)** Low-Frequency (LF) and **(b)** High-Frequency (HF) bands of the spectrum in the REST (left) and HUT (right) conditions. The strength of the frequency-specific causal couplings within the network is quantified in natural units ([nats]) of information. The different width of the arrows displays results from surrogate data analysis: absence of an arrow indicates low significance (*s <* 50%), a thin arrow indicates medium significance (50% ≤ *s <* 75%), while a thick arrow indicates high significance (*s* ≥ 75%).

As regards the coupled interactions between respiration and heart period, in the supine rest both LF and HF oscillations are captured by the MB causality measures (*s >* 75%, Fig. 5a,b, left). Moreover, higher values are detected along the direction *H* → *R* than *R* → *H* in both frequency bands. In HUT, HF is dominant and significance in LF is decreased along both directions (50% *< s <* 75%, Fig. 5a, right). In both conditions, the cardiovascular BR interaction is prevalent in LF, as indicated by the observed higher values and significance (Fig. 5, left panels: *s >* 50% in LF compared to *s <* 50% in HF during REST - the latter not shown; right panels: *s >* 75% in LF compared to 50% *< s <* 75% in HF during HUT). The opposite mechanism is visible in both bands with relatively low significance in REST (50% *< s <* 75%, Fig. 5, left panels). This direction is characterized by high significance in HF during HUT (Fig. 5b, right). Looking at the closed-loop interactions between arterial pressure and compliance, in both conditions the directed effect of MAP on AC is prevalent in LF, as indicated by the observed higher significance (50% *< s <* 75% in REST and *s >* 75% in HUT, Fig. 5a, against *s <* 50% in both conditions in HF, Fig. 5b). On the other hand, the directed effect of compliance on arterial pressure, i.e., *C* → *M*, has high significance during HUT (Fig. 5a,b, right) and during REST in HF (Fig. 5b, left), while medium significance in REST in LF (Fig. 5a, left). In the case of respiratory-vascular interactions, the closed-loop between respiration and mean BP shows higher significance in HUT than in REST along both directions and in both bands (*s >* 75% for *R* → *M* and 50% *< s <* 75% for *M* → *R*, Fig. 5a,b, right). The direction *R* → *M* shows medium significance in HF during REST, while it is not significantly relevant in LF (Fig. 5a,b, left). Additionally, respiration seems to have a role in driving arterial compliance beat-to-beat variability during the supine resting state and the orthostatic challenge, as demonstrated by a medium statistical significance (with the exception of *s <* 50% during REST in LF). However, this direct interaction is characterized by low values of the coupling in both frequency bands. Along the opposite direction, i.e., *C* → *R*, a weak interaction is detected during the resting state (Fig. 5a,b, left). Finally, the frequency-specific causal interaction from heart period to arterial compliance is visible with low coupling values and medium significance during HUT only in HF (Fig. 5b, right).

### D. Comparison between the supine resting state and the head-up tilt experimental conditions

In this section, we display results relevant to Fig. 6, with the aim to compare the REST and HUT experimental conditions.

**Figure 6.**
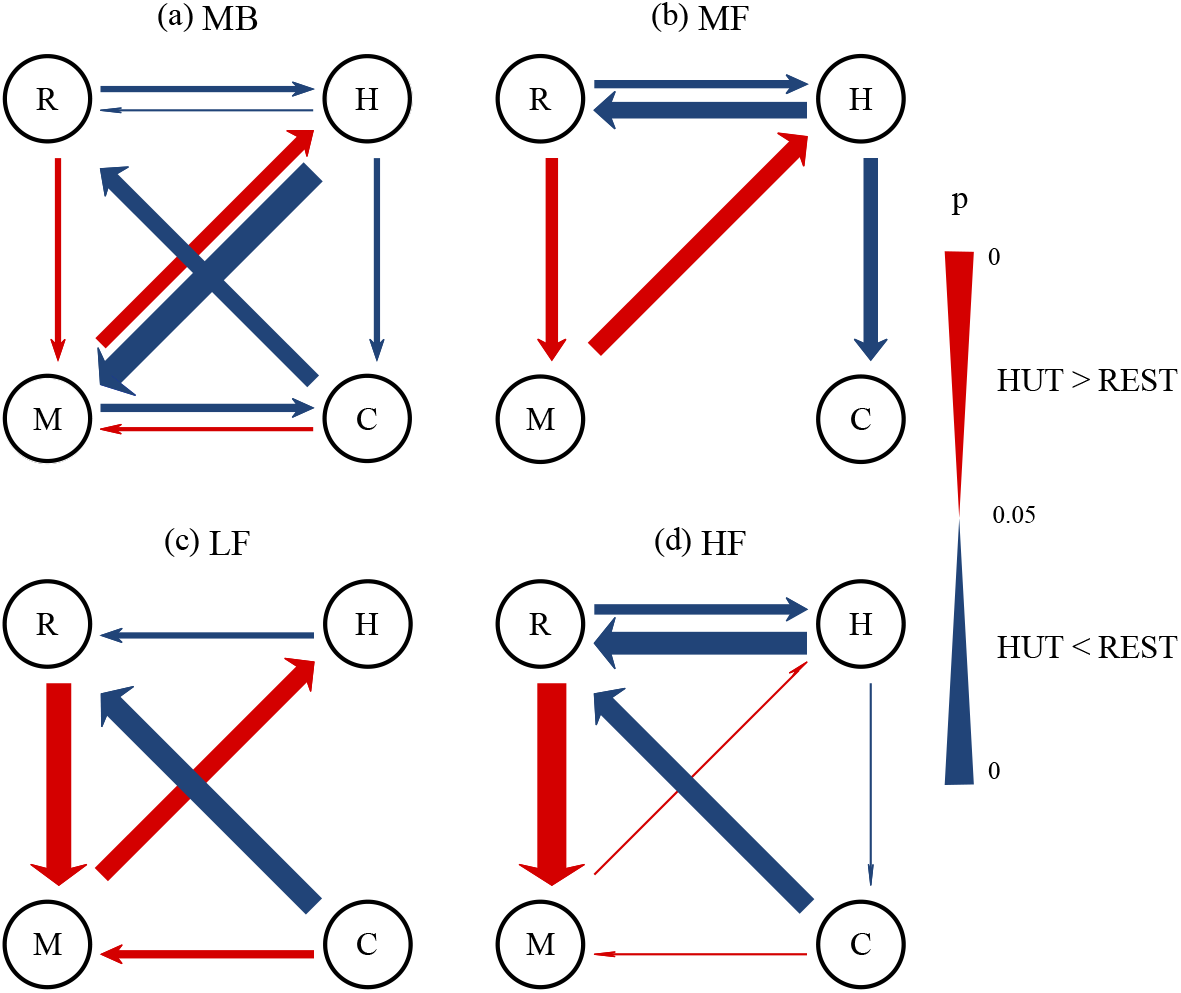
Directional networks indicating the significant differences between conditional causality measures computed in the REST and HUT conditions through the **(a)** model-based (MB) and **(b)** model-free (MF) approaches as well as in the **(c)** Low-Frequency (LF) and **(d)** High-Frequency (HF) bands of the spectrum. Wilcoxon signed rank test for paired data (*p <* 0.05): red, HUT *>* REST and blue, HUT *<* REST. Progressive change of the width of the arrows indicates increase/decrease of *p*-values in logarithmic scale. Arrows are shown only if the corresponding percentage of significance is above 50% for at least one of the conditions.

Cardiorespiratory interactions are strongly influenced by postural changes. The causal interaction *H* → *R* decreases significantly with HUT, and this diminished effect is visible exploiting both MB and MF approaches (Fig. 6a,b), with higher statistical relevance in HF than LF as indicated by the lower *p*-value (Fig. 6c,d). As regards the opposite direction, we detect a decrease of these dynamics with both MF and MB (only in the HF band) approaches (Fig. 6a,b,d). The BR effect of *M* on *H* increases significantly with HUT; the augmentation is captured by both methods and is mainly visible in the LF band of the spectrum (Fig. 6c). Conversely, the time domain MB causality measure detects a significant decrease of the feedforward interaction in HUT with a very low *p*-value (Fig. 6a). The MB method is able to detect an increase of the causal interaction *C* → *M*, with slightly higher statistical relevance in LF (Fig. 6c,d), and a decrease of *M* → *C* only in the time domain (Fig. 6a). Respiratory-related changes of MAP are enhanced in HUT, with *R* → *M* increasing using both approaches (Fig. 6a,b), and being relevantly visible in both HF and LF bands of the spectrum (Fig. 6c,d). The MB approach is also able to catch a decreased influence of the arterial compliance dynamics on the ventilatory activity in both bands of the spectrum and with significantly high *p*-value (Fig. 6a,c,d). Finally, the causal effect of heart period on arterial compliance decreases significantly during HUT for both approaches, with higher statistical importance by using MF methods (Fig. 6a,b). The parametric approach also reveals a significant difference between the two conditions in the HF band of the spectrum, even though this is very close to the significance threshold (Fig. 6d).

Fig. 7 shows the results of surrogate data analysis performed on the conditional TE measures computed using both model-based and model-free approaches (panels a,b), and considering their content in the low- and high-frequency bands of the spectrum (panels c,d) in both REST (top row) and HUT (bottom row) conditions. Heatmaps provide insightful suggestions on the statistical relevance that specific links acquire within the network, useful to identify the strongest connections in the supine rest and HUT conditions, as well as on their modification after the postural stress. Indeed, statistically significant differences in the conditional TE measures induced by the postural challenge are reflected by similar variations in the number of subjects for whom that measure is significant, i.e., an increase in the strength of the causal coupling is associated with a rise in its statistical relevance and viceversa. In addition, lower *p*-values reflecting the difference between REST and HUT conditions and higher differences in the number of subjects with statistically significant causality measures are often found in combination. Remarkably, some links within the network which undergo changes with HUT are characterized by a high and consistent importance in terms of the number of significant subjects in both conditions.

**Figure 7.**
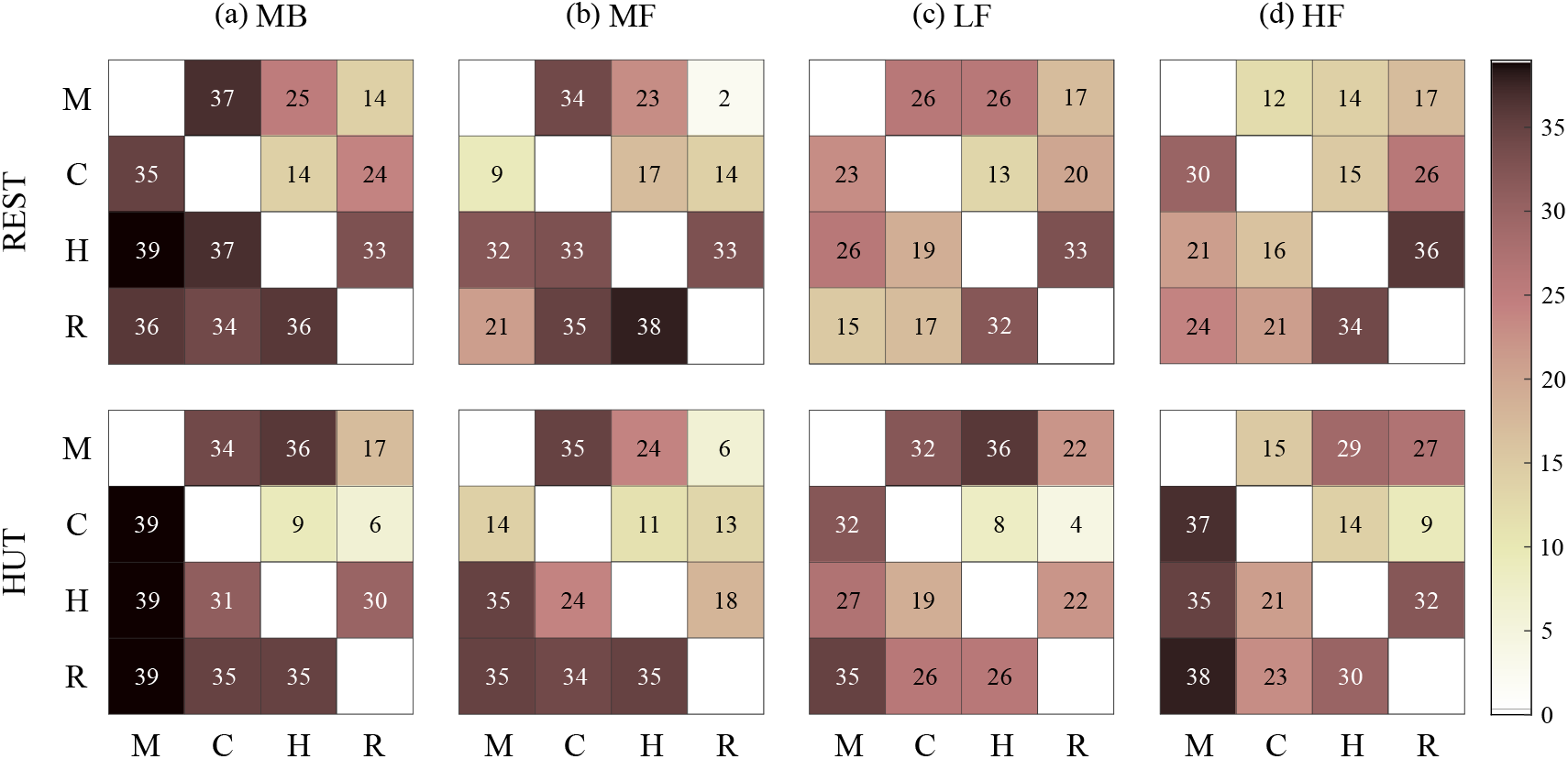
Heatmaps depicting the number of subjects for which surrogate data analysis provided statistically significant conditional TE measures computed by using the **(a)** model-based (MB) and **(b)** model-free (MF) approaches, as well as in the **(c)** Low-Frequency (LF) and **(d)** High-Frequency (HF) bands of the spectrum in the REST (top row) and HUT (bottom row) conditions. In each plot, source and target nodes are reported along rows and columns, respectively.

## III. DISCUSSION

This work points out the complex interplay between oscillations of cardiovascular variables in different experimental conditions, by considering the causal dependencies among pairs of time series within a dense and intricate network of interactions. In particular, the combination of model-based and model-free approaches, as well as of time domain and spectral measures of causality, allowed us to investigate both the linear and nonlinear characters of these coupled interactions, and to obtain frequency-resolved information on their oscillatory nature. From a physiological point of view, our findings reveal putative nonlinear mechanisms of interaction at the basis of some causal links within the analyzed network, and provide insights on low and high frequency patterns characterizing the interplay between physiological variables during the supine resting state and the orthostatic challenge.

### A. Methodological focus

In this work, AR model-based and nonlinear model-free approaches have been used to describe the interplay among the multiple nodes of complex biological networks in different physiological conditions. The comprehension of their advantages and disadvantages, as well as of their methodological differences, is crucial for understanding and interpreting these coupled interactions.

Theoretical definitions of TE, based on the concept of Shannon entropy [50], involve the use of variables of infinite dimensionality that sample the entire temporal evolution of the analysed processes [51]. In practice, however, realizations of these variables are of finite length. Dealing with such variables is simpler when working with linear models. Indeed, assuming Gaussianity, the interaction between the present and past states of the processes can be described by a vector AR model, and the information-based measures of TE can be computed from the covariance matrices of the processes [30]. MB approaches are sufficient for identifying the wide range of oscillatory interactions that take place under the hypothesis of linearity [52]. They allow the frequency domain representation of these multiple interactions, a feature which is extremely helpful in the study of physiological signals that usually exhibit oscillatory components in well-known frequency bands [53]. Moreover, linear AR models yields improved resolution, smoother spectra and can be applied to short segments of data [52].

However, model-free approaches are often necessary to overcome the drawback of losing generality typical of MB methods. Indeed, constraining the analysis on a specific model structure can limit the ability to detect and describe complex interactions and dynamic behaviors. For instance, the linear AR model fully captures the entropy variations of a Gaussian process, but is unable to identify potential nonlinear dynamic interactions [2]. MF approaches have the advantage of relaxing the linearity assumption, but they require direct, nonparametric estimation of probability distributions. Although its capacity of computing TE measures has been previously demonstrated [54], this method is complex and computationally demanding, and presents increasing bias as the dimensionality of multidimensional spaces spanned by the observed variables increases when working with short time series [45].

In Fig. 4, the strength and the statistical relevance of the causal interactions within the network are represented by different colors and widths of the arrows, respectively. The investigation of these two indicators, assessed via MB or MF approaches, may indicate prevalence of linear or nonlinear mechanisms beyond the observed phenomena. Indeed, despite the absolute values of the computed MB and MF measures are different as nonparametric estimators are known to underestimate causality [28], it is still important to compare the networks obtained using the two approaches in terms of presence/absence of specific links and their statistical weight.

On the other hand, time domain analysis of causality provides an overall evaluation of the network dynamics, without resolving frequency-specific information typical of physiological processes. The utilization of spectral conditional TE measures is necessary when dealing with rhythmic processes rich of oscillatory content, often localized in different bands of the frequency spectrum.

In Fig. 5, the different colors and widths of the arrows allow to identify the strength and statistical relevance of the frequency-specific causal interactions within the network. The comparison between causality measures assessed in different bands of the spectrum is essential to characterize low and high frequency dynamic patterns of the observed processes. Noteworthy, this spectral representation is limited to the lack of the zero-lag effect modelling, as specified in Sect. I D 1 and Sect. I D 3. For this reason, it is not possible to directly relate the frequency domain with the time domain results obtained with the MB approach. This acquires a meaning especially in Fig. 6, where the absence of a specific arrow in the spectral domain (panels c,d) and its presence in the time domain (panel a) can be indicative of tilt-induced relevant changes of instantaneous causal effects, such as in the case of the decreased feedforward mechanism.

### B. The network physiology behind the resting state

The study of resting state cardiovascular networks is of great interest for research and clinical applications. Physiological parameters fluctuate in order to maintain body homeostasis, reflecting the ability of healthy subjects to respond appropriately to physiological changes [55]. In this section, we interpret results relevant to the supine resting state by describing the mechanisms underlying the beat-to-beat variabilities of heart rate, blood pressure and arterial compliance within the investigated network of interactions.

#### 1. Heart rate variability

It is well known that heart rate is one of the physiological parameters characterized by the highest variability. Heart rate variability (HRV) varies with age and gender [55–57], and its lack or depression have been described as indicators of several pathological states, e.g., nervous system disorders [58], diabetes [59], arterial hypertension [60], and myocardial infarction [61]. The physiology behind the regulation of cardiac dynamics is complex, but most studies agree that the main components of the normal sinus rhythm are related to the control exerted by the autonomic nervous system (ANS) [55, 57, 62]. During ventilation, the activity of the sinoatrial (SA) node is directly influenced by the modulation of vagal neurons directed to the heart, controlled by the central respiratory drives (direct communication between respiratory and cardiomotor centers), the lung inflation reflex and the changes in arterial blood pressure transferred to heart rate via baroreflex [19, 55, 63–65]. These mechanisms result in the so-called respiratory sinus arrhytmia (RSA), for which there is an increase of heart rate during the inspiration phase and a decrease during the expiration phase of ventilation [19, 62, 66]. Recent studies documented the underexplored complexity of cardiorespiratory interactions, highlighting the important role exerted by synchronization mechanisms such as cardioventilatory coupling [62, 67]. Specifically, the latter is the entrainment phenomenon in which ventilation and heartbeats become synchronized in whole number ratios and heartbeats fall in constant timing relationship with the onset of inspiration [5, 68, 69]. It has been suggested that the cardioventilatory coupling, whose relationship with RSA needs further investigation, is likely caused by a cardiac-induced onset of inspiration through an unknown haemodynamic afferent pathway [68]. It is thought that the positioning of heartbeats where they are maximally affected by RSA could have some physiological benefit in improving cardiopulmonary performance; indeed, it has been found to enhance efficiency in oxygen delivery by matching ventilation and perfusion [68, 69].

In this work, we aimed to disentangle and discuss these intricate physiological matters. Both the MB and MF estimation approaches were able to detect significantly relevant causal interactions along the directions *R* → *H* and *H* → *R* (Fig. 4a,b, left, and Fig. 7a,b, top), suggesting that the mechanisms involved in the regulation of cardiorespiratory dynamics are significantly strong and bidirectional. Remarkably, some studies documented the importance of using nonlinear approaches, either based on entropy measures or nonlinear AR models, to investigate the complex interactions between respiration and heart rate [6, 48, 70]. Indeed, it was demonstrated that the interplay between the cardiac and respiratory systems may lead to the rise of nonlinear RSA dynamics [6, 70], herein well identified by the MF estimator (Fig. 4b, left, and Fig. 7b, top). Moreover, as also highlighted in [71], cardiorespiratory patterns of interaction are characterized by a relevant instantaneous influence of the breathing dynamics on the cardiac activity. In our work, this effect was accounted for, as detailed in Sect. I D 3. Remarkably, while it was set a priori given physiological knowledge in the case of the MB method [26], for the MF conditional causality measure along the *R* → *H* interaction the zero-lag sample of the RESP series was selected for more than 50% of subjects, thus confirming the major role played by respiratory-driven instantaneous interactions directed to the heart. On the other hand, the spectral MB approach, despite not modelling these zero-lag effects, was still able to evidence the synchronicity of cardiac and respiratory oscillations in the breathing band, i.e., the HF band (Fig. 5b, left, and Fig. 7d, top), with major strength along the direction *H* → *R*. Noteworthy, while the physiological causes and effects of the RSA mechanism (i.e., the causal coupling *R* → *H*) have been widely studied and discussed throughout the past decades [19, 66, 72], very little is known about the cardiac-driven respiratory variability. It is widely believed that the coupling between the cardiovascular and respiratory systems is unidirectional, i.e., the respiratory rhythm influences the heart rate via vagal nerve traffic oscillations and, to a much lower extent, via direct mechanical influence on the sinoatrial node [73]. Nevertheless, some evidence conflicts with this theory suggesting that the respiratory oscillator in the central nervous system is not always dominant, i.e., the cardiorespiratory coupling can be bidirectional in some specific cases. Moreover, a physiological model to study cardiorespiratory coupling showed that both the influence of respiration on heartbeat and the influence of heartbeat on respiration are important for cardiorespiratory synchronization [74]. Herein, we attempt to confirm this hypothesis and suggest the presence of strong closed-loop interactions involving cardiorespiratory dynamics separate from the confounding effects of BP and arterial compliance as a result of the utilization of conditional causality measures. Although we do not delve into physiological details as regards the interaction directed from the heart to respiratory variability, past studies have explored the effect of mechanisms such as baroreceptor reflexes, as well as the role of the autonomic nervous system, in mediating these cardiorespiratory interactions [5].

The role of HRV should be considered when discussing the complex interplay between cardiovascular variables. We found a weak but significant link along the baroreflex, i.e., from MAP to RR, using both estimators (Fig. 4a,b, left, and Fig. 7a,b, top), mainly characterized by low-frequency oscillatory dynamics (Fig. 5a, left, and Fig. 7c, top). This is a well-established result, widely investigated in the literature, and related to the presence of LF oscillatory rhythms in the variability of RR and BP [75]. Remarkably, the significant predominance of LF oscillations along the baroreflex was already pointed out in several previous studies [76, 77].

#### 2. Blood pressure variability

In the supine resting state, together with the heart rate, blood pressure also exhibits spontaneous fluctuations and is strongly influenced by ventilatory and sympathetic activities [78]. Remarkably, although the study of physiological mechanisms involving BP variability (BPV) is usually assessed by SAP or DAP measurements, the use of mean arterial pressure has recently gained support [35]. It was found that, especially at rest, the mean BP is most tightly coupled with RR [35], thus opening the possibility to exploit this input signal instead of systolic BP for BR analysis. Starting from this frontier, in this work we investigated MAP-based cardiovascular and vascular-respiratory interactions.

With regard to the influence exerted by respiration on BP, the significant causal interaction from RESP to MAP (Fig. 4a,b, left) is likely due to the cyclic intrathoracic pressure changes that occur during ventilation and lead to alterations in venous return and fluctuations in BP values [79], independently of heart rate changes inducing cardiac filling fluctuations. This effect, even though with low values of the causal coupling, has higher significance in the respiratory band (Fig. 5b, left, and Fig. 7d, top). Overall, our results indicate that the respiratory-driven variability of MAP during the resting state may show a linear character, due to both the observed higher values and significance of the directed coupling *R* → *M* assessed through the time domain MB approach compared to the MF approach (Fig. 4a,b, left, and Fig. 7a,b, top).

Besides, the resting-state BPV is known to be strongly influenced by a relevant non-baroreflex effect (commonly known as feedforward) led by changes in heart rate, reasonably due to the Windkessel [80] and/or Frank–Starling [81] mechanisms. In this regard, some studies documented a predominance of the cardiovascular effects of heart rate on diastolic BP (run-off effect interaction), as well as of diastolic on systolic BP, which together constitute the pathway heart rate - diastolic BP - systolic BP explaining the high amount of information flowing from RR to SAP in the supine resting state [31, 82]. Despite these mechanisms have not been investigated nor analysed in terms of the MAP time series yet, the strong dependence of MAP on DAP and SAP values (with MAP being computed from the integral of the BP curve between two consecutive DAP values) makes the MAP series interesting from this perspective as well. Indeed, our results confirm a strong feedforward mechanism and thus highlight the major role played by MAP within this chain of interactions (Fig. 4a,b, left, and Fig. 7a,b, top). Remarkably, the selection of a percentage of zero-lag samples by the MF method slightly lower than 50% across subjects, as well as the (i) highly significant causal interaction *H* → *M* detected through the time domain MB (Fig. 4a, left) together with (ii) the lower significance of the same link in the LF and HF bands (Fig. 5a,b, left), suggest that the instantaneous effect directed from heart period to mean BP may have an important role in characterizing cardiovascular dynamics. Moreover, spectral and surrogate data analyses showed that this interaction is mainly confined to the LF band of the spectrum in the resting state (Fig. 5a,b, left, and Fig. 7c,d, top).

Causal effects on BP can also be ascribed to arterial compliance variability. The directed interaction from AC to MAP appears to be significantly relevant in the case of causality measures computed with the MB approach (Fig. 4a, left, and Fig. 7a, top). Although these dynamics seem to be more evident in pathological conditions or with increasing age [83, 84], the beat-to-beat modulation of systolic BP due to arterial compliance variability may play an important role even in youth and healthy physiological conditions. As suggested by previous studies, this can be explained by considering the potential effects of the SV modulations (thus of the compliance) on the arterial system (thus on BP variability) conveyed by respiration [85] and venous blood pool control [86]. Moreover, previous studies documented an important role of PVR as a source of MAP changes [87], although the causal interaction *PV R* → *BP* has been investigated only with bivariate approaches (see, e.g., [32]). It has been suggested that PVR, determined by vasomotion in smaller arteries, influences BP by diastolic blood pressure decay (short-term changes) and changes in venous return associated with blood redistribution (long-term changes). Since AC is herein computed from *τ* and PVR, we can argue that the influence of AC on MAP can be due to the direct effect of PVR on BP.

#### 3. Arterial compliance variability

With regard to the variability of arterial compliance, the investigation of this parameter has emerged recently in the literature. Recent studies have observed how its regulatory mechanisms are complex and not clearly understood yet [9, 10, 12]. Remarkably, being a non negligible physiological factor strongly influencing AC, the direct action of sympathetic nerves on the vascular tone of large vessels may play the role of an unobserved confounder when interpreting the coupled interactions from and to AC [88].

As supported by previous studies [11], blood pressure plays an important role in the modulation of pulse wave velocity (PWV) in the supine rest. Assuming PWV as a surrogate index of compliance [9], this is likely reflected by a significant causal interaction along the direction *M* → *C*, revealed by both MB and MF approaches (Fig. 4a,b, left, and Fig. 7a,b, top) and mainly located in the LF band of the spectrum (Fig. 5a, left, and Fig. 7c, top). It is worth noting that the relationship between mean arterial pressure and compliance can be affected by several factors. Indeed, both the weight of the zero-lag effect of MAP on AC, which was selected in more than 50% of subjects by the MF estimator, and the sympathetic activity of the ANS, which can be considered as an LF-driven common source for BP and AC, may have an influence on the strength and the significance of the coupling directed from mean BP to vessel compliance. Additionally, the sympathetic-driven effect of cardiac contractility along the cardiac inotropic arm of the baroreflex should be also taken into account [32], since beat-to-beat modifications of BP induced by internal or external stimuli at rest may have an impact on the relationship between BP and AC.

Our results also reveal the influence of both heart period and breathing patterns on arterial compliance at rest. Several studies have demonstrated a significant effect of heart rate fluctuations on compliance variability independently of autonomic control and BPV [11, 84, 89, 90]. An increase in heart rate has been associated to modifications of the cardiac cycle and a reduction in the time available for recoil, resulting in vascular stiffening and arterial compliance decrease, due to the viscous and inertial components of the arterial walls [84, 91]. Remarkably, the relationship between heart rate and vascular compliance has been investigated indirectly in the most of previous works on the topic. Indeed, assuming PWV as a surrogate for AC [9], some studies have shown that PWV is indirectly related to RR independently of BP [11, 84, 89, 90], supporting the hypothesis that the effect of heart period on arterial compliance could show nonlinear features in the supine resting state. Despite the sympathetic activity of the ANS has an effect directed to the heart which cannot be neglected and may play a common drive role here, our results show a significant strong directed interaction from RR to AC which cannot be ascribed to the confounding effects of respiration nor BP variability (Fig. 4a,b, left, and Fig. 7a,b, top). Furthermore, we found that the zero-lag effect from RR to AC is selected in more that 70% of subjects in the resting state using the MF approach, demonstrating the importance of embedding instantaneous effects into the analysis. Noteworthy, MB spectral values of *H* → *C* suggest that this interaction falls into the low-significance range (i.e., *s <* 50%), proving that this method fails in detecting significant patterns of frequency-specific causality (Fig. 5a,b, left, and Fig. 7c,d, top). To support this finding, we remind that the proposed spectral measures suffer from the lack of the zero-lag interaction modelling: neglecting this effect may lead to underestimate causality in the frequency domain.

As regards the influence exerted by respiratory variability on compliance, it is known that small changes of intrapleural or intrathoracic pressure and lung volume during spontaneous ventilation can independently affect arterial compliance through changes in atrial filling or preload and modifications in transmural pressure exerted on the arterial wall [92]. This can determine modifications of some cardiovascular parameters, such as SV, as well as of the mechanical properties of the arterial wall [93, 94]. Looking at our results, the directed interaction *R* → *C* was detected with the highest significance (*s >* 75%) by both approaches (Fig. 4a,b, left, and Fig. 7a,b, top), and was found to be mostly located in the HF band of the spectrum with 50% *< s <* 75% (Fig 5b, left, and Fig. 7d, top). Interestingly, despite respiratory activity may have an indirect influence on compliance through modulation of heart period and BP, herein we get around the issue by utilizing conditional causality measures and separating the link from confounding external effects. This confirms that a direct effect of respiratory variability on vessel compliance is likely to occur in the supine resting state and may reflect the effect of cyclic changes in transmural pressure on arterial wall stiffness. Moreover, the high significance of this causal link in both time domain MB and MF measures can be ascribed to a great influence of the zero-lag effect, which indeed was selected for more than 50% of subjects by the MF non-uniform embedding procedure.

### C. Modification of network interdependencies in response to the orthostatic stress

In this section, we focus on how the causal relationships explored in the supine resting state are adjusted in response to the orthostatic challenge, in order to better understand the complex physiology behind the tilt-induced modification of the interactions among the investigated variables.

The orthostatic challenge is known to alter important physiological mechanisms operating in the resting state condition. Orthostasis has been associated to venous pooling of the blood in the lower portion of the body, thus decreasing cardiac filling, cardiac output and stroke volume [95]; in turn, this determines a drop of arterial blood pressure sensed by baroreceptors, vagal inhibition and sympathetic activation directed to the heart and vessels [13, 14]. As previously observed in literature [18, 32], these mechanisms lead to a reduction of HRV and to an increase of mean BPV (Fig. 3a,b, bottom row). Moreover, we observe that the tilt-induced decrease of mean AC (Fig. 3c, top row) is associated with a decrease of the average inter-beat interval (Fig. 3a, top row), confirmed in the literature [11], as well as with a decrease of MAP (Fig. 3b, top row). Assuming that the mean and systolic BP behave similarly in response to the orthostatic challenge [35], our results are in accordance with a previous study [10], where a shift of the SAP-AC curve towards lower values of compliance and systolic BP during HUT was detected. The observation of decreased AC, together with the well-known shift of the SAP-AC curve, reflects a change in the arterial tree properties occurring with orthostasis and thus has been associated with vasomotor activity [10, 96]. Indeed, the diminished values of AC during HUT may be attributed to the tilt-induced sympathetic activation, leading to smooth muscle cells constriction in elastic arteries. Contrarily, the decrease of arterial compliance variability is a novel observation (Fig. 3c, bottom row). Since we are not aware of any relevant citation for this phenomenon, we can speculate about the potential underlying mechanisms: (i) decreased sympathetic control oscillations directed to elastic vessels, even though in contrast to increased oscillations in sympathetic control to resistance vessels (mostly arterioles); influence of decreased HRV (Fig. 3a, bottom row), and (iii) lower influence of MAP and RR on stiffer arteries during sympathetic activation.

#### 1. Cardiorespiratory patterns during head-up tilt

As depicted in Fig. 3d and confirmed in previous studies [97, 98], the tilt-induced decreased respiratory rates and increased tidal volumes, together with the the well-known parasympathetic activity withdrawal during HUT, leads to a significant RSA weakening, here identified by both MB and MF approaches (Fig. 4, and Fig. 6a,b,d). This is confirmed by previous studies [99, 100] and has been associated to a dominance of the direct central mechanism mediating RSA (i.e., direct communication between respiratory and cardiomotor centers) at supine rest and an increased indirect peripheral mechanism (i.e., ventilation-associated BP changes transferred to heart rate via baroreflex) during orthostasis [19, 63–65]. Remarkably, the latter is clearly reflected in our results by the tilt-induced augmented causal interactions *R* → *M* and *M* → *H* (Fig. 6), which together constitute the indirect peripheral mechanism controlling RSA and will be discussed in the following subsection.

A reduction of the cardiorespiratory coupling is also detected by looking at the causal direction *H* → *R* and more visible with the MF approach (Fig. 4b, and Fig. 6b). This decrease has major statistical relevance using MB measures in the HF band intended as lower *p*-value (Fig. 5b, and Fig. 6d), despite the number of significant measures remains high across conditions and its variation from REST to HUT is thus negligible (Fig. 7d). In this regard, it is worth noting that the highest variation in the number of significant measures is found with the MF approach (Fig. 7b), which indeed may better capture the tilt-induced modification of this cardiorespiratory mechanism. Nevertheless, very little is known about the cardiac-driven respiratory variability and its modification with tilt, as already pointed out in Sect III B. We learn from literature that, although this coupling is poorly seen in alert and active humans, it was clearly observed during relaxation, sleep, anaesthesia and, generally, in conditions of low cognitive and behavioural activity [68]. Here, the significant decrease of the causal coupling along the direction *H* → *R* may be associated to the gradually decrease of the complex cardiorespiratory interactions during situations of sympathetic activation [65].

#### 2. Involvement of blood pressure variability in tilt-induced physiological responses

The increased influence of breathing activity on mean arterial pressure variability, revealed by the MF approach (Fig. 4b and Fig. 6b) and by MB approach in both frequency bands (Fig. 4a, Fig. 5, and Fig. 6a,c,d), is representative of ventilation-associated BP changes occurring with tilt and due to alterations in intrathoracic pressure, instantaneous lung volume, venous return to the heart and cardiac output [101]. Previous studies [7, 102, 103] suggested that respiratory-related fluctuations of BP can be ascribed to both the mechanical thoracic coupling between respiration and vasculature and the effects of respiratory induced fluctuations of RR. While RSA is thought to buffer respiratory-related BP oscillations during the supine rest [7, 103], the mechanical influences of respiration on arterial pressure are thought to be greater in the upright than the supine position. We attempt to confirm this hypothesis: since the causal effect of respiration on mean BP is conditioned on the knowledge of all the other variables in the network, the observed increase of the directed coupling *R* → *M* is solely due to the enhancement of the mechanical influences of respiration on arterial pressure. Moreover, it was demonstrated that these effects are present at the low frequencies but predominate at frequencies higher than 0.25 Hz [102, 104], as confirmed here (the variation of the causal coupling *R* → *M* with tilt shows slightly higher statistical relevance in HF than LF). Noteworthy, the significant increase of this effect is supported by an expected rise in the number of significant measures detected with surrogate data analysis (Fig. 7c,d).

In the complex regulation of BP variability during postural stress, arterial compliance is also involved. Specifically, we found a significant increase of the causal interaction *C* → *M* by using the MB approach and in both frequency bands (Fig. 4a, Fig. 5, and Fig. 6a,c,d). As far as we know, the present study represents the first attempt to investigate the causal interactions within a network of four physiological processes comprising arterial compliance. Previous works on the topic suggested that an effect of vascular stiffness on arterial pressure may be related to the LF rhythms conveyed to the arterial system through SV modulations; the latter are likely to depend on respiration [85] and on venous blood pool control [86]. Nevertheless, we are not aware of previous studies which made an effort to explain the physiological mechanisms behind the modification of these causal interconnections during orthostasis. This is not a straightforward job, but we believe that our findings may have an important role in future works which aim to disentangle the complex interplay between BP and vessel properties. At the level of the current knowledge, we suggest that a relevant portion of the tilt-induced changes of mean BP should be ascribed to the short-term arterial compliance variability.

It was previously shown how a transient change in carotid arterial compliance is responsible for a sudden change in the cardiac baroreflex response [88]. In this regard, the relationship BP-AC may have effects on the interactions between heart period and BP. Moreover, several works argue that the tilt-induced significantly higher awareness of baroreflex control, due to blood redistribution and baroreceptor unloading during orthostasis, can be also related to the confounding effect of respiration [18, 31, 82]. In this study, since any external driver of BP variability, i.e., the compliance and respiration variability, is ruled out by the conditioning procedure, we suggest that the observed enhancement of the baroreflex control with the importance of the gravitational stimulus (Fig. 6) is only due to the closed-loop interplay between heart period and arterial pressure, thus confirming previous results obtained in older studies on the topic [31, 32, 65, 76, 77]. Noteworthy, the significant relevance of modifications along the baroreflex is likely related to the increase of LF power in the variability of RR and BP [75–77] (Fig. 5a and Fig. 6c). The feedforward direction of the cardiovascular closed-loop, i.e., *H* → *M*, is affected by the postural stress, as detected by the MB approach (Fig. 4a, and Fig. 6a). The highly statistically relevant decrease of this causal coupling is not associated with a correspondent diminished number of significant measures detected with surrogate data analysis (Fig. 7a), meaning that the strength of the interaction remains considerably noteworthy within the network. This result is in line with previous studies [31], where the effect of HUT was found to be prominent for the directions *RR* → *DAP* and *DAP* → *SAP*. The authors suggested that these results can reflect the involvement of other mechanisms influencing DAP and/or contributing to the strength of systolic contraction, such as changes in the PVR associated with tilt-induced sympathetic activation and sympathetic nervous system influence on cardiac contractility, respectively. In our study, despite we cannot rule out the role exerted by the sympathetic branch of the ANS, we can argue that some other putatively involved factors herein investigated, such as changes in vessel compliance associated with sympathetic activation and alterations in breathing patterns, may have significant effects on the strength of the feedforward interaction and act as confounders for this link. However, with the utilization of conditional causality measures, we have the ability to eliminate these common causes. All these findings support the idea that, during tilt, changes of BP variability are related to the increase of sympathetic activity induced by the postural stress [82], as well as to the decreased buffering of BP mediated by HRV.

#### 3. Regulation of arterial compliance variability during postural stress

Here, we speculate about the mechanisms involved in the beat-to-beat regulation of arterial compliance during head-up tilt. Interestingly, we observe a significant decrease of the causal interaction *M* → *C* using the MB approach (Fig. 4a and Fig. 6a), thus suggesting that orthostasis affects the strength of this link probably due to stiffer vessels. This is a novel finding, since a thorough inspection of this interplay, which is not a straightforward task, is still lacking in the literature. Similar investigations have been carried out in the past years, where the relationship between systolic BP and compliance was found to hold for a given phase but shift during the HUT phase towards lower values of systolic BP and compliance [10, 105]. However, the present work goes beyond the classical bivariate analysis, with the aim of ruling out the confounding effects of cardiac and respiratory variables. Following this rationale, we speculate that the decrease of *µ*_*C*_ in HUT (Fig. 3c, top row) can be partly MAP-dependent, i.e., due to a diminished transfer of information from BP. Nevertheless, it has been suggested that it could depend also on other mechanisms such as sympathetic nervous system activation and/or changes in PVR during HUT [96]. Noteworthy, the diminished causal interaction *M* → *C* appears statistically significant by using the MB approach but not the frequency-specific measures in the LF and HF bands (Fig. 5), probably due to the important role of the zero-lag effect of BP on compliance not embedded in spectral analysis. As a proof, although the MF approach did not detect this decrease, it included zero-lag samples of MAP for about 50% of subjects along this direction, thus confirming the remarkable weigh of the instantaneous interaction when taking into account the relationship between arterial pressure and compliance. A decreased coupling along the direction *H* → *C* is also observed with both MB and MF approaches (Fig. 4, and Fig. 6a,b), as confirmed by surrogate data analysis (Fig. 7a,b), but is not frequency-specific (in HF, the statistical relevance is minimal, Fig. 6d). This finding can be related to the important influence of zero-lag causality (more than half subjects if investigated through the MF approach) on the observed interaction, which could reflect pure mechanical visco-elastic effect. Again, these considerations are absolutely novel and need further studies on the topic to provide deeper physiological reasoning beyond the observed phenomena.

Arterial compliance may play an important role in driving respiratory variability during postural stress, as suggested by the decrease of the causal coupling *C* → *R* detected with the MB approach (Fig. 6a, and Fig. 7a) and visible in both bands of the spectrum (Fig. 5, Fig. 6c,d, and Fig. 7c,d). We argue that these dynamics exhibit predominantly linear characteristics and speculate that they may be related to the mechanical interaction between the arterial system and ventilation.

### D. Limitations and future studies

Several caveats affecting the physiological interpretation of our results can be drawn from this study.

First, it is well known that changes affecting the variability of driver processes can have significant repercussions on the computed causality measures. Tilt-induced diminished variabilities, such as in the case of HRV, may be partly responsible for the observed decrease of some causal interactions, as happens for the directed links *H* → *R, H* → *M* or *H* → *C* (Fig. 6). This effect cannot be ruled out but it should be taken into account when translating the obtained results into physiological considerations.

Moreover, in spite of the effort made to build a network with great physiological resonance, some nodes are still missing which may play a significant role as confounders of the investigated interactions. As an example, it is worth mentioning the remarkable role of vasomotion, i.e., vasoconstriction and vasodilation phenomena due to sympathetic activity directed to the vessels, which may have an influence on arterial compliance [96, 106]. These considerations, together with the insights we provided in the discussion of our results, lead to the conclusion that the sympathetic activity represents a common source to compliance, HRV, PVR and BP. This represents a valuable limitation in the study of physiological dynamics, as the sympathetic firing cannot be measured nor disregarded but undoubtedly has significant effects on the interplay among variables in the considered network.

We believe that our work has the potential to be groundbreaking in the field of network inference as regards cardiovascular and cardiorespiratory dynamics. However, we are aware of the methodological limitations of the applied approach in accounting for the intricate dynamics of these systems. Indeed, even though we consider here the multivariate nature of the investigated network, the use of conditional causality measures has the main drawback of not taking into account the high-order effects of complex control mechanisms. It has been demonstrated that both pairwise and fully conditioned TE analyses may encounter challenges in the presence of synergy or redundancy in time series data; specifically, fully conditioned causality may fail to reveal redundant effects among the observed processes [107]. Furthermore, understanding how the multiple structural links functionally interact in determining the complex dynamics of the whole network may shed light on the intricate regulation mechanisms leading to homeostatic balance. To this end, further analysis, based either on decomposing the information shared into unique, synergistic, and redundant contributions [108] or exploring the redundant/synergistic role of network links in information processing, should be carried on to achieve a more detailed and comprehensive description of the node-, link- and network-wise behaviors of the considered physiological network [25, 109].

## CONCLUSION

This work shed light on the complex dynamics involved in the short-term cardiovascular regulation during the resting state and in response to the postural stress. The integration of arterial compliance variability within a four-node network including heart period, arterial pressure and respiration allowed a more complete description of cardiovascular and cardiorespiratory mechanisms. This approach favored the inference of the physiological network behind the homeostatic control of cardiac, vascular and respiratory activities, showing the ability of healthy subjects to respond adequately to postural stress and the associated modifications. Results confirmed already known physiological control mechanisms involving cardiovascular dynamics, e.g., the BR loading, or cardiac and respiratory activities, e.g., the RSA mechanism, but also led to interpret less studied mechanisms involving, e.g., the influence of cardiac and blood pressure dynamics on arterial compliance, as well as the effect of compliance on breathing patterns.

Furthermore, the thorough application of the directional causality measure of conditional transfer entropy using both linear parametric and nonlinear model-free approaches, as well as the focus on frequency-specific dynamics, allows for a comprehensive description of the physiological mechanisms involved in the resting state and during the postural stress. Among the several physiological interactions identified, our analyses show that cardiorespiratory interactions represent the strongest mechanism within the investigated network, and how this interplay is characterized by high-frequency oscillatory content involving nonlinear dynamics with a relevant zero-lag interaction effect. Furthermore, besides the well-known cardiac-respiratory influence on tilt-induced changes of blood pressure, our results evidenced the emergence of a linear directed effect of arterial compliance on mean arterial pressure dynamics. Indeed, for the first time we assessed the important role of arterial compliance within a complex network of cardiovascular variables, evidencing that short-term mechanisms affecting the beat-to-beat variability of this parameter are essentially within-beat and nonlinear. The latter is a novel observation which poses the basis for further and deeper investigation.

## APPENDIX

### COMPUTATION OF CONDITIONAL TRANSFER ENTROPY MEASURES

Let us consider a network of *M* dynamic systems, whose activity is described by the set of *M* random processes **X** = {*X*_1_, …, *X*_*M*_}. The information-theoretic measure of conditional transfer entropy (TE) [36] can be exploited to quantify the direct causal links among the nodes of the network, as it allows the evaluation of the directed information flow from a source process *X*_*i*_ to a target process *X*_*j*_ (*i, j* = 1, …, *M, i* ≠ *j*) accounting for the effect of the remaining *M* − 2 processes collected in **Z** = **X** \{*X*_*i*_, *X*_*j*_}. Specifically, considering the random variables describing the present and past states of the processes *X*_*i*_, *X*_*j*_ and **Z** at time *n*, i.e., *X*_*j,n*_, *X*_*i,n*_, **Z**_*n*_, and 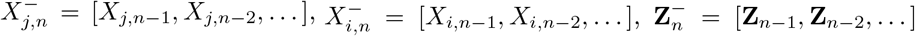, respectively, the conditional TE is formulated as [36]:

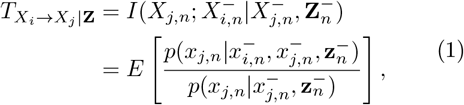

where *E* [·] is the expectation operator, *p*(·|·) the conditional probability, and *I*(·;·|·) the conditional mutual information.

### A.1 Model-based formulation of conditional causality

Assuming that the activity of the investigated network can be described by means of linear Gaussian stationary processes, an AR model-based (MB) definition of conditional TE can be exploited to describe the causal relationships between the observed data.

For Gaussian processes, it was shown that the concept of TE, formulated as a measure of causal information transfer between joint processes [37], is entirely equivalent to the concept of Wiener-Granger causality (GC) [42], which was formalized in terms of linear autoregression by Granger [38]. A straightforward time domain formulation of conditional GC based on a multivariate linear AR model and extended to include zero-lag effects has been proposed in [26]. This measure is used in this study to characterize the time domain causal relationships among the multiple nodes of the investigated physiological network. Conversely, frequency domain MB formulations of conditional GC have been introduced in [29], and lately discussed and applied in [30, 39]. Nevertheless, these frequency-specific measures are defined in the context of strictly causal AR models, i.e., models where zero-lag interactions are neglected. Despite a number of methods have been proposed to overcome this issue in terms of pairwise causality between two (blocks of) time series, especially in the time domain [26, 46, 47], spectral conditional causality measures still lack a clear formulation in the presence of zero-lag effects. For this reason, in this study we used spectral measures of conditional GC as in [29], without accounting for the effects due to instantaneous interactions among the multiple nodes of the network.

Following this rationale, in this work conditional GC was assessed in its traditional formulation based on vector autoregressive (VAR) models [38, 39]. Specifically, the dynamic interactions among the *M* processes in **X** can be described by the VAR model [43]:

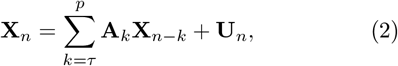

where *p* is the model order, defining the maximum lag used to quantify interactions, *τ* is the starting lag, **X**_*n*_ = [*X*_1,*n*_, …, *X*_*M,n*_]^*T*^ is a *M* -dimensional vector collecting the present state of all processes, **X**_*n*−*k*_ = [*X*_1,*n*−*k*_, …, *X*_*M,n*−*k*_]^⊺^ is a *M* -dimensional column vector collecting the past state of all processes at lag *k*, **A**_*k*_ is the *M* × *M* matrix of the model coefficients relating the present with the past of the processes at lag *k*, and **U**_*n*_ = [*U*_1,*n*_, …, *U*_*M,n*_]^*T*^ is a *M* -dimensional vector with *M* × *M* positive definite covariance matrix **Σ**, collecting the present state of the zero-mean white noises {*U*_1_, …, *U*_*M*_}, the latter uncorrelated with **X**_*n*−*k*_, *k* = *τ*, …, *p*.

To compute measures of GC from *X*_*i*_ to *X*_*j*_ conditioned on **Z**, a restricted linear regression is derived starting from the VAR representation in (2):

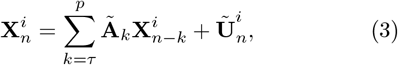

Where 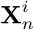 and 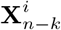 are (*M* − 1)-dimensional vectors collecting the present and past states of all processes in **X** except *X*_*i*_, i.e., **X**^*i*^ = {*X*_1_, …, *X*_*i*−1_, *X*_*i*+1_, …, *X*_*M*_}, Ã_*k*_ is the (*M* − 1) × (*M* − 1) matrix of model coefficients relating the present and past samples of the processes in **X**^*i*^, and 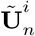 is a (*M* −1)-dimensional vector with covariance matrix **Λ**, collecting the present state of the zero-mean white noises {*Ũ*_1_, …, *Ũ*_*i*−1_, *Ũ*_*i*+1_, …, *Ũ*_*M*_}.

#### Extended and strictly causal VAR models

The initial lag *τ* should be set appropriately for computing conditional GC measures. Specifically, when *τ* = 0, zero-lag interactions are taken into account in the description of the relationship between the present and past states of the *M* processes; the VAR models (2), (3) are referred to as *extended* [46], from which measures of conditional causality in the time domain can be derived [26] (Sect. A.1.1). Conversely, setting *τ* = 1 means that instantaneous effects are discarded. The VAR models (2), (3) are referred to as *strictly causal* [46], from which measures of conditional causality in the frequency domain can be derived [29, 30] (Sect. A.1.2).

##### A.1.1 Time domain formulation

To compute extended measures of conditional GC from *X*_*i*_ to *X*_*j*_, unrestricted and restricted linear regressions combining both instantaneous and lagged effects (*τ* = 0) are derived from the VAR models in (2) and (3):

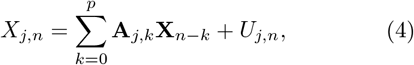

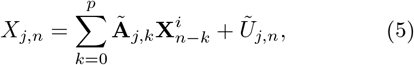

where **A**_*j,k*_ and **Ã** _*j,k*_ represent the *j*^th^-row coefficients of **A**_*k*_ and **Ã** _*k*_, respectively, weighting the present and past samples of all processes in **X** for the unrestricted regression (4), and the present and past samples of all processes in **X**^*i*^ for the restricted regression (5). The residuals of the unrestricted (4) and restricted (5) regressions have variances 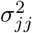 and 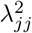, respectively, and are extracted as the *jj*^*th*^ elements of the covariance matrices **Σ** and **Λ**, respectively. The extended conditional GC measure is computed by comparing the variance of the residuals of the two models,

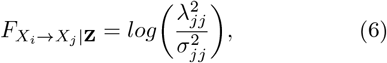

and, in the case of Gaussian processes, it corresponds to the 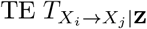 (1) up to a factor 2, i.e., 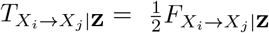 [30, 110].

##### A.1.2 Frequency domain formulation

The VAR models (2) and (3) provide global representations of the overall multivariate process **X** and its subset **X**^*i*^, respectively, and can be exploited to compute conditional causality measures in the frequency domain. To this end, here we set *τ* = 1 and thus neglect instantaneous effects between the processes.

The Fourier Transforms (FTs) of (2) and (3) are taken to derive, respectively:

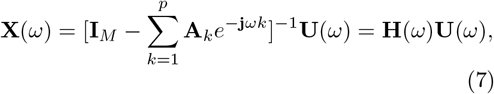

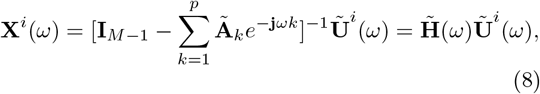

where **X**(*ω*), **X**^*i*^(*ω*), **U**(*ω*), and **Ũ** ^*i*^(*ω*) are the FTs of **X**_*n*_, 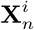, **U**_*n*_, and **Ũ** _*n*_, respectively; *ω* ∈ [−*π, π*] is the normalized angular frequency (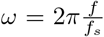 with 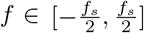, being *f*_*s*_ the sampling frequency), 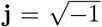, and **I**_*M*_ and **I**_*M*−1_ are the *M* -dimensional and (*M* − 1)- dimensional identity matrices, respectively. The *M* × *M* matrix **H**(*ω*) contains the transfer functions relating the FTs of the innovation processes in **U** to the FTs of the processes in **X**. The (*M* −1) ×(*M* −1) matrix 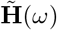 contains the transfer functions relating the FTs of the innovation processes in **Ũ** to the FTs of the processes in **X**^*i*^.

Following the procedure described in [29, 30], it is possible to combine (7) and (8) to derive measures of conditional causality. Specifically, considering the transfer function **Q**(*ω*) = **G**(*ω*)**H**(*ω*), with **G**(*ω*) being a *M* × *M* matrix computed rearranging the elements of 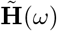 as in [29, 30], it is possible to write the power spectral density (PSD) of a *M* -dimensional vector process comprising **X** and the innovations of the VAR model in (3) as:

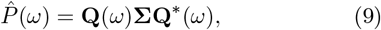

where * stands for conjugate transpose. Taking the element of 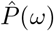 associated with *Ũ*_*j*_, i.e., 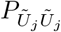, the conditional GC from *X*_*i*_ to *X*_*j*_, defined as the portion of the power spectrum associated with *U*_*j*_, can be computed as [29, 30]:

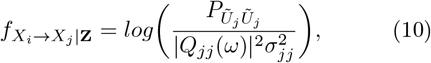

where *Q*_*jj*_(*ω*) is the *jj*^*th*^ element of **Q**(*ω*). The measure in (10) is non-negative at each frequency *ω*; its integration along well-defined spectral ranges returns the corresponding time domain conditional GC measures in the selected bands, thus fulfilling the so-called spectral integration property [30, 110, 111]. As for the measure defined in (6), it can be demonstrated that, under the assumption of gaussianity, the spectral conditional GC in (10) corresponds to the spectral 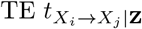 up to a factor 2, i.e., 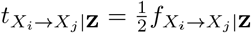 [30, 110].

### A.2 Model-free formulation of conditional causality

The nonparametric approach used in this work for evaluating direct causal measures is based on the idea that the probability density around a data point is inversely related to the distance from its nearest samples, i.e., on the *k*-nearest neighbour (KNN) method [40]. Although this approach is widely employed for its reliability and robustness, the accuracy of the estimates is lower when the estimated probability distributions are simply replaced in (1). Indeed, the combination of terms evaluated on spaces of different dimensions causes the presence of significant bias in the estimates. The formulation introduced by Kraskov, Stögbauer and Grassberger limits this bias by using the same range search in all the spaces after defining the searching distance in the highest dimensional space [41].

Exploiting the decomposition of the conditional mutual information in different entropy quantities, the conditional TE can be estimated as [28]:

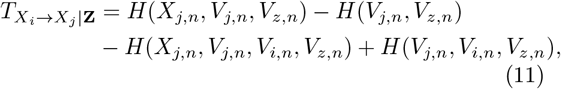

where *V*_*j,n*_, *V*_*i,n*_ and *V*_*z,n*_ are the embedding vectors of dimension *d*_*j*_, *d*_*i*_ and *d*_*z*_, respectively, approximating 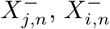, and 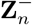.

Using the maximum norm to calculate distances, the entropy term in the highest dimensional space, i.e., [*X*_*j,n*_, *V*_*n*_] with *V*_*n*_ = [*V*_*j,n*_, *V*_*i,n*_, *V*_*z,n*_], is estimated as:

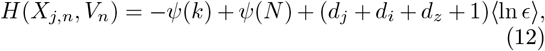

where ⟨·⟩ is the average operator, *ψ*(·) the digamma function, *N* the number of observations, and *ϵ/*2 the distance between the considered sample and its *k*^*th*^ neighbour. The entropy terms defined in the projected lower dimensional spaces are indeed estimated as:

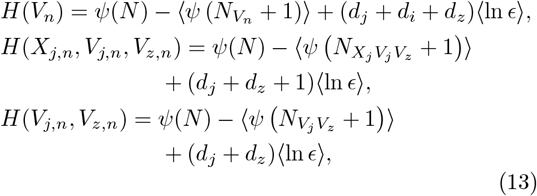

where 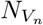, 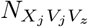 and 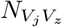 are the number of points at distances smaller than *ϵ/*2 from the considered sample in the spaces *V*_*n*_, [*X*_*j,n*_*V*_*j,n*_*V*_*z,n*_] and [*V*_*j,n*_*V*_*z,n*_], respectively.

Substituting the entropy terms (12,13) in (11), the nearest-neighbour estimate of the conditional TE is obtained as:

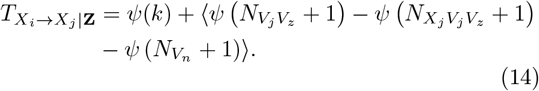

#### Embedding procedure

Finding embedding vectors that closely approximate the infinite-dimensional past states of the processes is a critical step in estimating theoretical-information measures using model-free (MF) approaches. When working with data of finite length, e.g., the 300 samples typically used for the analysis of short-term physiological time series, the employment of high-dimensional detailed vectors to provide a more complete description of past processes leads to the curse of dimensionality and unreliable estimates of entropy quantities [45]. An alternative selection technique to the uniform embedding approach, which simply uses a fixed number of equally spaced samples, was introduced to limit the size of the descriptive patterns and maximise their informational content about process dynamics. Specifically, the non-uniform embedding approach described in [28] was exploited in this work.

Considering a set of candidates including all possible states of the processes up to a maximum lag *L*, i.e., 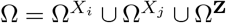 being 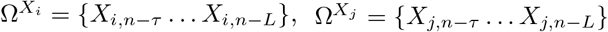 and Ω^Z^ = {**Z**_*n*−*τ*_ … **Z**_*n*−*L*_}with *τ* the starting lag, an iterative approach is applied to select the components of *V*_*n*_ which maximize the information shared by the target process *X*_*j,n*_ and the candidates. Starting with an empty embedding vector 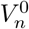, for steps *d* ≥ 1, the candidates Ŵ_*n*_ are selected from a subset 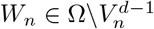 as:

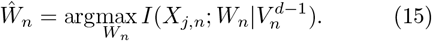

After each step, a surrogate-based approach is used to test the significance of the selected candidates. Specifically, the conditional mutual information 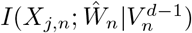 is compared with a threshold obtained as the 100(1 − *α*)^*th*^ percentile of a distribution resulted computing the same measure over *N*_*s*_ realizations of the process obtained shuffling randomly and independently the samples of Ŵ_*n*_ and *X*_*j,n*_. Thus, the selected candidate is added to the embedding vector if 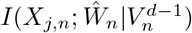 is higher than the threshold, resulting in 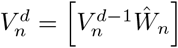.

In this work *N*_*s*_ = 100 realizations of surrogate data were generated and the significance threshold fixed to *α* = 0.05.

#### GRANTS

L.S. and L.F. were supported by PRIN 2022 project “HONEST-High-Order Dynamical Networks in Computational Neuroscience and Physiology: an Information-Theoretic Framework” (funded by MUR, code 2022YMHNPY, CUP B53D23003020006). M.J. was supported by grant VEGA no. 1/0107/25.

## AUTHOR CONTRIBUTIONS

C.B., L.S., L.F. and M.J. conceived and designed research; C.B. and L.S. performed experiments; C.B. and L.S. analyzed data; C.B., L.S., L.F. and M.J. interpreted results of experiments; C.B. and L.S. prepared figures; C.B. and L.S. drafted manuscript; C.B., L.S., L.F. and M.J. edited and revised manuscript; C.B., L.S., L.F. and M.J. approved final version of manuscript.

